# Automatic classification of behavior in zebrafish larvae

**DOI:** 10.1101/052324

**Authors:** Adrien Jouary, German Sumbre

## Abstract

Zebrafish larvae navigate the environment by discrete episode of propulsion called bouts. We introduce a novel method for automatically classifying tail bouts. A supervised soft-clustering algorithm to categorize tail bouts into 5 categories of movements: Scoot, Asymmetrical Scoot, Routine Turn, C Bend and Burst. Tail bouts were correctly classified with 82% chance while errors in the classification occurred mostly between similar categories. Although previous studies have performed categorization of behavior in free-swimming conditions, our method does not rely on the analysis of the larva’s trajectory and is thus compatible with both free-swimming and functional imaging in head-fixed condition.

## Introduction

The zebrafish larva propels itself through a sub-carangiform pattern of body undulations. The oscillations of the tail are coordinated with pectoral fins movements. At the larval stage, zebrafish locomotor patterns are characterized by swimming episodes intermingled with non-swimming episodes called “beat and glide”. The discrete segments formed by the beat and glide swim in larvae are called tail bout, the range of durations of tail bouts is 80-500 ms, the range of tail beat frequency is 30-100 Hz.^1^ The quantification of behaviors is therefore greatly facilitated because of the discrete nature of locomotion. Zebrafish larvae exhibit a variety of tail bouts: they include slow scoot (also called forward swim), routine turn, J turn or C bend illustrated in Table 1. These categories were described according to the tail movements as well as the kinematics of the trajectories.^2–4^ Because they are defined by the properties of the trajectory, this categorization is not suited for the head-fixed conditions, where the trajectories are unknown.

**Table 1.**
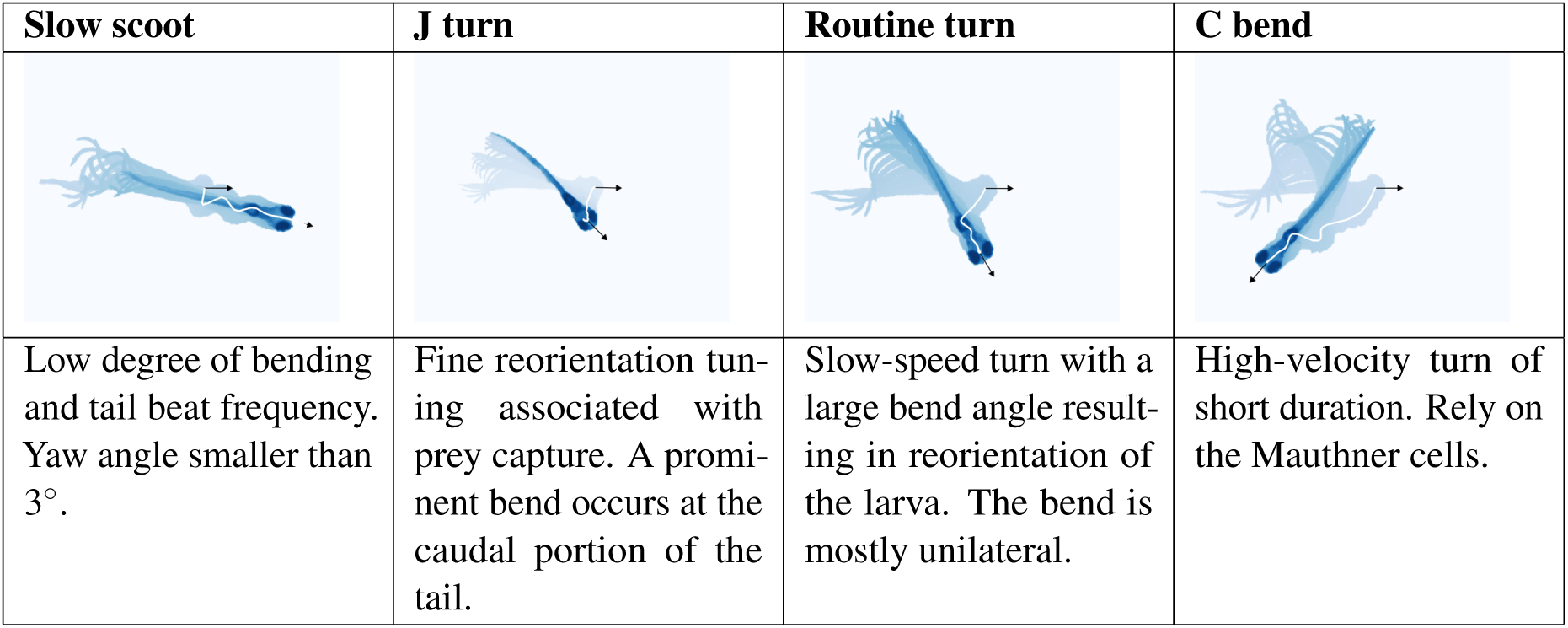
Stereotypical tail movements. Each column represents a typical tail movement and its characteristics. The middle column shows the superimposition of image of the larva during the tail bout. The trajectory of the head is shown by a white line, the black arrows represent the head orientation at the beginning and end of the bout.

| Slow scoot | J turn | Routine turn | C bend |
| --- | --- | --- | --- |
| Low degree of bending and tail beat frequency. Yaw angle smaller than $3^\circ$. | Fine reorientation tuning associated with prey capture. A prominent bend occurs at the caudal portion of the tail. | Slow-speed turn with a large bend angle resulting in reorientation of the larva. The bend is mostly unilateral. | High-velocity turn of short duration. Rely on the Mauthner cells. |

Here, we propose a method for classifying tail bouts, using dynamic time warping for comparing tail movements. The repertoire of tail bouts recorded in head-fixed larvae appeared to form a continuum rather than a series of discrete group of movement types. In order to obtain a parsimonious representation of this continuum of locomotor actions, we trained a supervised classifier to identify 5 different categories of movements: Scoot, Asymmetrical Scoot, Routine Turn, C Bend and Burst.

## Results

We recorded the behavior of 25 head-restrained larvae for a period of 4h. During the first hour, visual stimulations consisting of whole-field motion or dark flashes were presented in order to observe visually induced behavior. For the last 3h, a homogeneous non-patterned illumination was projected below the larva. This experiment enabled us to collect ~16000 individual bout movements.

## Quantification of tail movements

Quantifying locomotor action requires choosing an appropriate level for the movement’s description, between the most detailed analysis of muscle activations to the simple binary detection of movements. In order to build a classifier, we needed to estimate how similar or different two movements were. To that end, we computed both the curvature along the tail^5^ and the deflection index.^6^ In order to compute the curvature. To find the ‘skeleton’ of the larva’s body, we computed the barycenter of the larva’s image along a circle weighted by the pixel’s intensity. The first circle was centered to a position between the eyes. By iteration, the skeleton of the fish was obtained (Fig. 1.A). A cubic spline was fitted along the skeleton and the curvature was computed along this curve. Both techniques were implemented in real time in C++ in order to reduce the amount of post-processing analysis.

**Figure 1.**
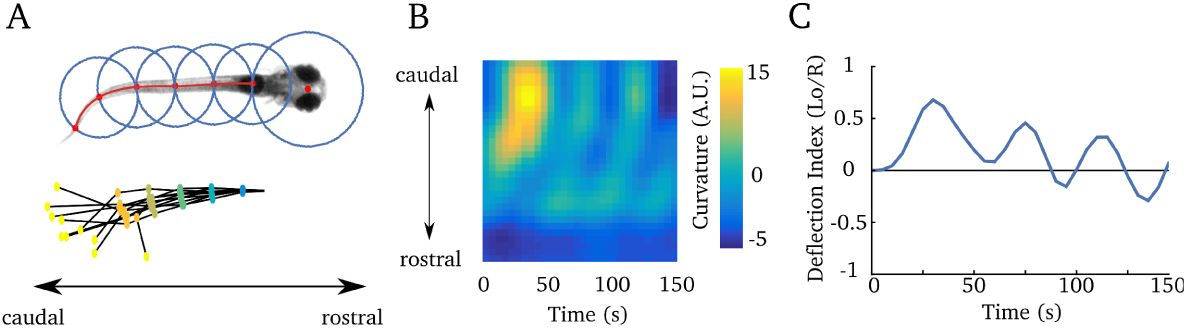
Quantification of tail bouts (**A**) Upper panel: A larva is overlaid on the circles used to compute the skeleton of the larva’s body. The first center was placed between the two eyes. Successive centers were computed as a barycenter of pixel intensity on the successive circles. Lower panel: example of the positions of the skeleton during a tail bout. Each skeleton represents a different time point during a bout. The color dots represent the center of the circles in (A). (**B**) Evolution of the tail curvature during a bout. At each time point, the curvature was computed according to a cubic spline fitted along the skeleton. (C) Deflection index along time. The same tail bout was used for the 3 panels.

## Similarity between tail movements

Using the curvature and deflection index, we established a measure of similarity between movements. A similarity measure was necessary to find categories of movements. We benchmarked two alternative measures: feature and distance based similarity.

The feature-based description of a tail movement added heterogeneous measurements: tail bending, amplitude and change in orientation. In head-fixed larva, the features can only use the time series of the tail curvature. Each bout was described by a high-dimensional vector of these features. Its dimensionality was then reduced using principal component analysis (see Supplementary methods for details and Fig. 2.A).

**Figure 2.**
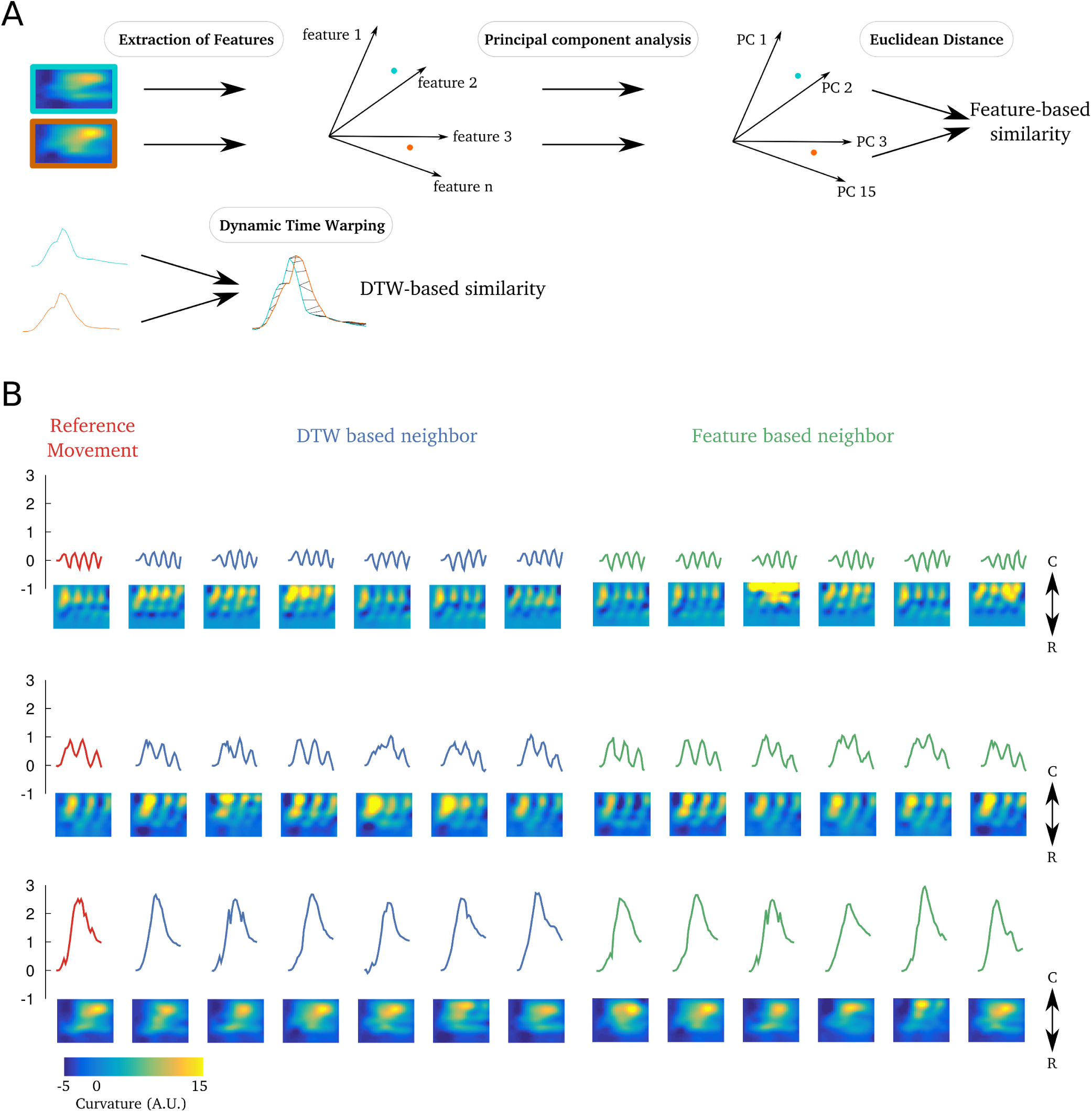
Comparison of Feature and DTW-based similarity measurements. (A) Scheme of the procedure used to compute the feature and DTW similarity measurements. Upper panel: feature vectors were extracted from the curvature matrix using the procedure described in Supp Methods, after a projection of the feature vectors on the 15 principal components with the highest variance. The euclidean distance in the principal components’ subspace was finally computed. Lower panel: the dynamic time warping was computed from the deflection index of the tail using the algorithm described in Supplementary methods. (**B**) Each rows shows the nearest neighbor of three reference movements (in red) computed using features-based distance (in blue) and DTW similarity (in green). In each row, individual tail bouts were represented using the tail deflection index and the corresponding tail curvature (below). Arrows indicat the caudo-rostral axis. The colorbar is common to all curvatures.

The distance-based approach uses Dynamic Time Warping (DTW) to compare the time series of tail deflection. DTW is a similarity measure between time series that is robust to small deformation of the time series. In contrast, with euclidean distance, the main advantages of DTW is that it recognized similar shapes, even when the time series present signal transformation such as shifting or scaling (see Supplementary methods for details and Fig.2.A).

Fig. 2.B shows the nearest neighbors using both methods for three reference tail bouts. Despite the fact that the DTW neighbors were computed using the deflection index alone, it was sufficient to recover neighbors that shared similar curvatures. The distance obtained from features of the curvature and the similarity based on DTW of the deflection index had a correlation of 0.77. Thus, we chose to use DTW to measure similarity because it is more direct and does not rely on an arbitrary choice of the different bout’s features. we then performed cluster analysis in order to define different movement categories.

## Classification of tail movements

The goal of clusterization is to find groups of movements that are more similar with each other than to those in other groups. Each tail deflection index was duplicated to its opposite in order to obtain a symmetric library of movements. Fig. 3.B shows the t-distributed stochastic neighbor embedding (t-SNE) of all tail movements recorded. This method embedded high-dimensional data into a two dimensional representation where each point corresponds to a tail movement (see Supplementary Methods for details). The axis in the t-SNE embedding should not be interpreted, but the local distances between points in 2D reflect the similarity between the movements that they represent. Thus, neighboring points correspond to similar tail movements.

**Figure 3.**
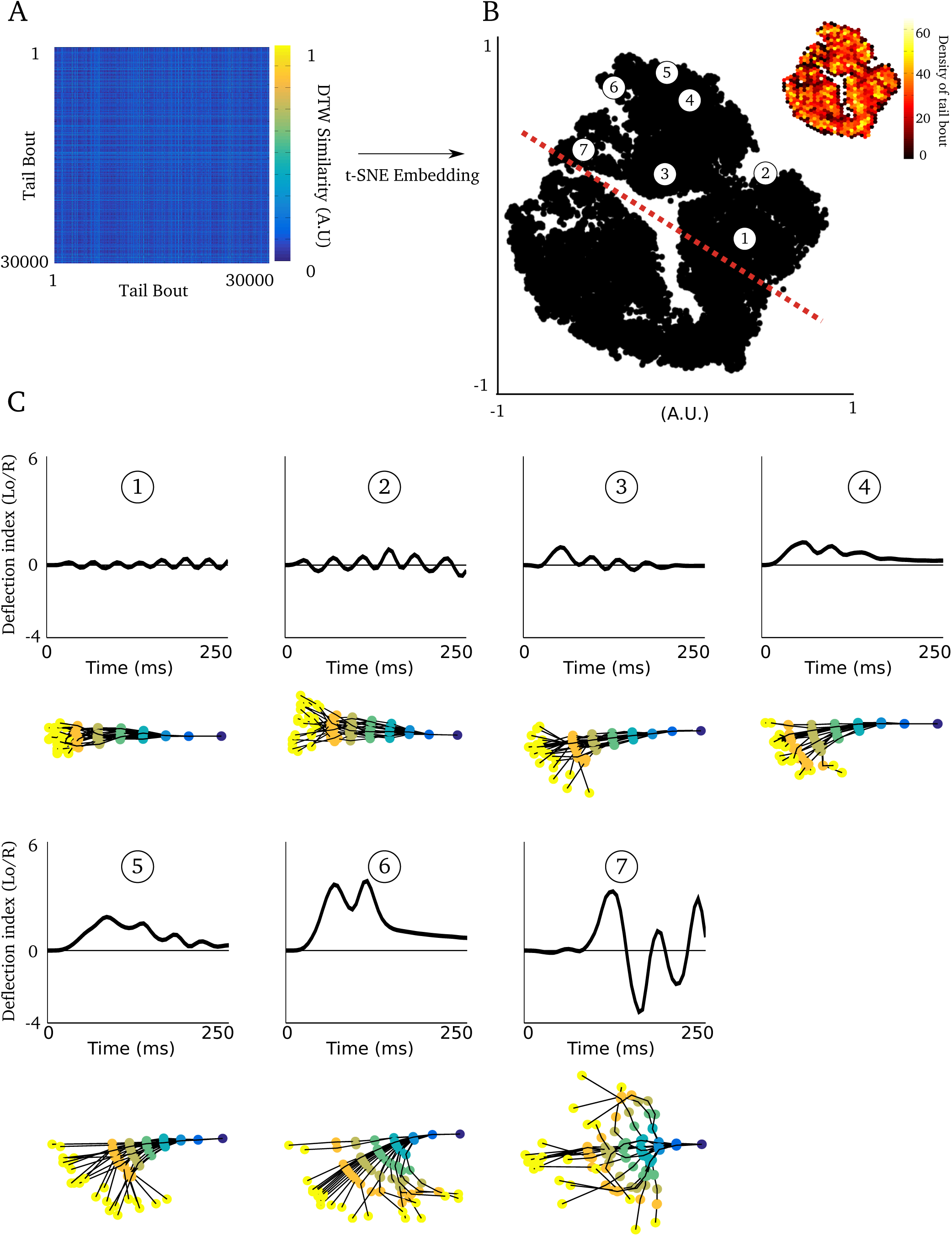
Continuum of tail kinematics. (**A**) Distance matrix computed on the library of evoked and spontaneous tail bouts. (B) t-SNE embedding of the distance matrix. Each dot represents a tail movement. The red dashed line indicates the approximate position of the axis corresponding to the symmetry between movements with opposite tail deflection indexes. Inset shows the density computed in hexagonal bins. Note that the density is homogeneous along all regions of the graph, suggesting that in head-restrained conditions, the repertoire of the larva’s tail movements can not be classified in discrete movement types, but is rather follows a continuum. (**C**) Illustration of the continuum between tail movements for a selection of 7 different tail bouts, a symmetric version could be found in the other side of the dashed axis. The deflection index (top) as well as the corresponding skeleton (bottom) are displayed. The position of the tail bouts in the t-SNE embedding is shown by an indexed circle.

This visualization did not reveal clearly isolated groups of points or local maxima in density, but rather a continuum between tail movements (Fig. 3.B,C). Unsupervised classification methods, such as k-mean or density based clustering^7^ isolated a cluster formed by forward scoot, but failed to find the other categories of movement. Therefore, We used supervised clustering to define categories.

Supervised techniques infer the category of a movement using a set of manually labeled movements.^8^ We chose 8 categories: Scoot (symmetric), Asymmetric Scoot (positive or negative), Routine Turn (positive or negative), C Bend (positive or negative) and Burst (symmetric). J Turns (Table 1.) were associated with Asymmetric Scoots or Routine Turns depending on their amplitude. Escapes were grouped along with bursts.

In order to account for the continuity between categories, we used a soft-clustering algorithm: the fuzzy K-nearest neighbor (see Supplementary Methods for details). Using soft-clustering, a movement can belong to more than one cluster, thus it is defined according to a set of membership levels. For each movement, the membership level associated with a category can be interpreted as the probability that the movement belongs to this category. The category of a movement was attributed according to the highest membership value.

This method resulted in an accuracy in classification of the movements of 82% estimated using cross-validation. Cross-validation consisted in iteratively removing the manually labeled tail bouts of one larva (the test set) and comparing its label to the label inferred from the remaining library of tail bouts (the training set). Fig. 4.A shows the misclassification matrix, errors in classification occurred mostly between neighboring clusters, e.g. scoot movements were unlikely to be classified as a burst. Moreover, the maximal value of the membership was lower for misclassified movements than for well classified ones (Fig. 4.B). Thus, errors in classification occurred mostly when there was an ambiguity between the membership levels. We then colored movements in the t-SNE according to their membership, the geometry of the 2D embedding was nicely represented by the membership (Fig. 4.C). It should be noted than the representation obtained by t-SNE was not used for the classification but only for visualization purpose. The classification relied only on the distance matrix obtained using DTW (Fig. 4.A).

**Figure 4.**
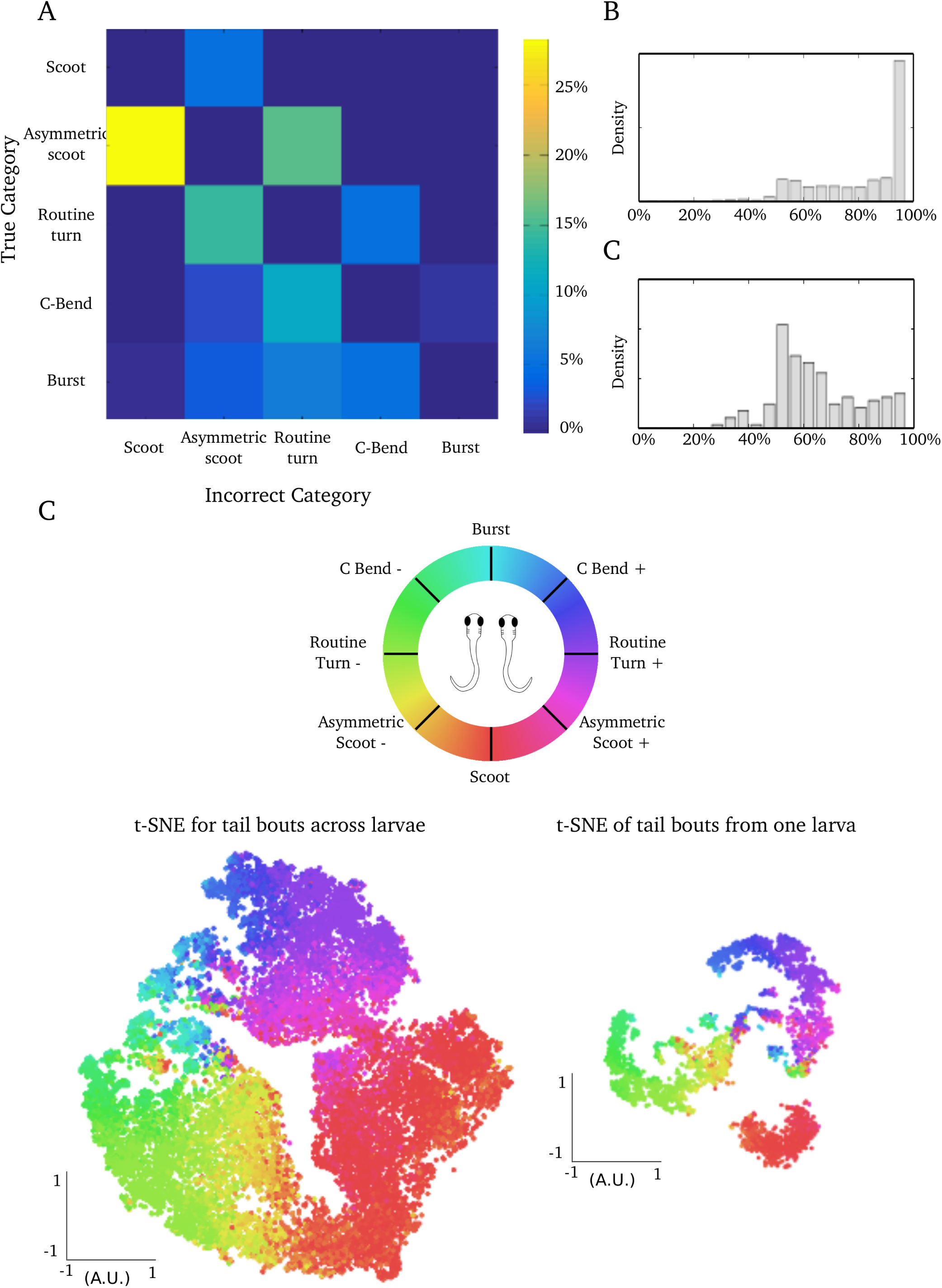
Classification of tail bouts. (**A**) The misclassification matrix is a contingency table of the error in classification obtained during cross-validation. Rows represent the true category (defined by user) and columns represent the categories that were erroneously attributed during cross-validation. High values are centered around the diagonal indicating that error occurred primarily between neighboring categories, especially between scoots and asymmetric scoots. (**B**) Upper panel: Distribution of the maximum level of membership for movements in the test set. Lower panel: Distribution of the maximum level of membership for movements in the test set that were misclassified. Errors in classification were associated with more uniform level of membership suggesting ambiguity. (**C**) Movements in the t-SNE embedding graph were color-coded according to their membership to the different categories. Each of the 8 category of movements was associated with an angle (top). For each movement, the average angle weighted by its membership values was computed and associated with the hue of its color (shown in the circular HSV colormap). The convention for the laterality is indicated by the larva’s scheme corresponding to positive or negative deflection indexes. Bottom left: t-SNE computed from the entire library of tail bouts. Bottom right: t-SNE computed for only one larva. The axis in both embedding graphs were different but displayed similar structures.

We used cluster analysis in order to get a parsimonious description of locomotor actions. The categories chosen are known to represent the action of distinct group of neurons or different behavioral context. In contrast with previous studies (,^24^), this method does not rely on the trajectory of the larva and can therefore be applied to study movements performed in conditions compatible with brain imaging.

## Distribution of movements according to evoked or spontaneous conditions

In order to confirm that the choice of categories was relevant to study the behavioral repertoire, we analyzed how stimulus biased the distribution of movements across categories (Fig. 5). Compared to spontaneous movements, whole-field motion in the caudo-rostral direction induced a large majority of scoot movements. In comparison, the proportion of scoot movements was smaller when the grid moved in the rostro-caudal direction. Finally, dark flashes induced a proportion of C bends 2.5 times higher than during spontaneous behavors, consistent with previous reports.^9^

**Figure 5.**
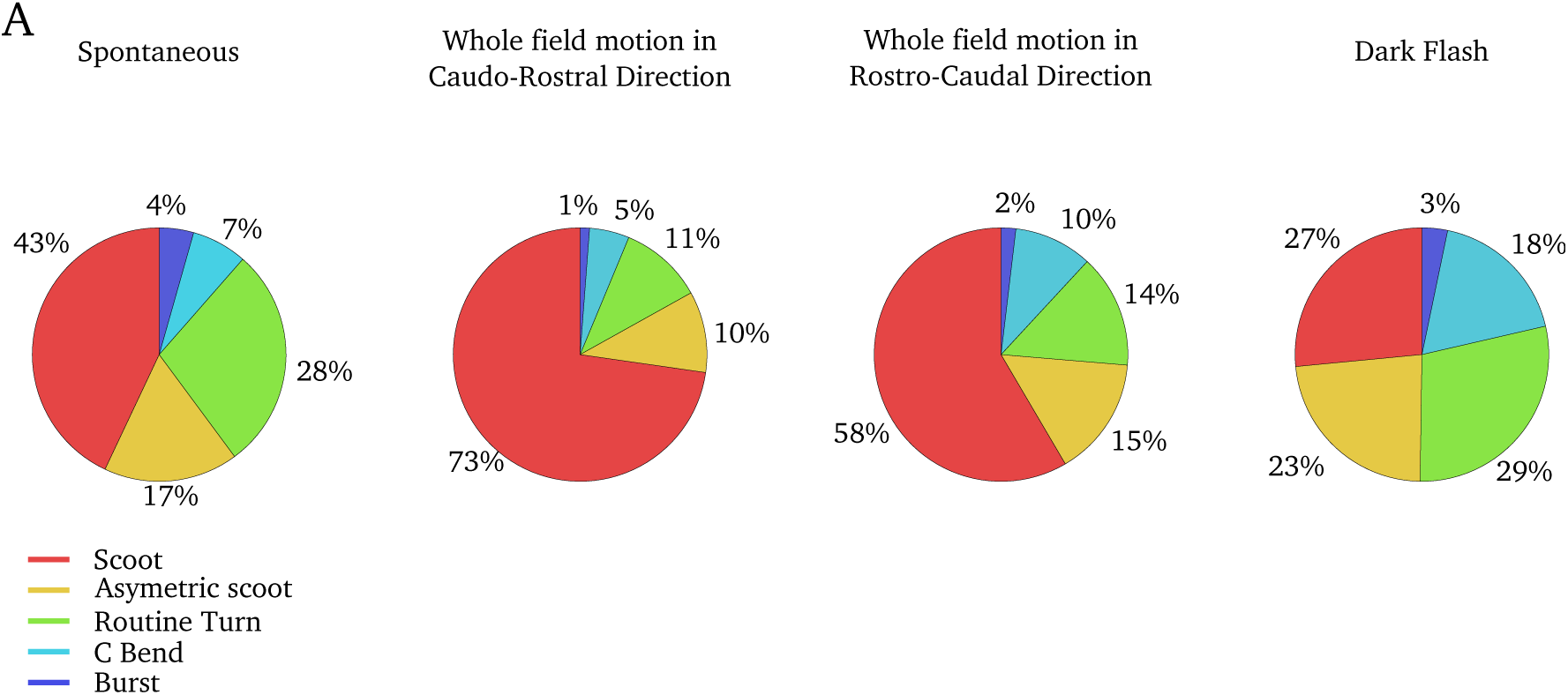
Distribution of movements in induced and spontaneous conditions. Each column shows a pie chart of the distribution between categories of movements in spontaneous or sensory evoked conditions.

## Discussion

The natural segmentation of movement events associated with a reasonably stereotyped locomotor repertoire is ideally suited for large-scale characterization of zebrafish’s ethogram.

We found that the repertoire of tail bouts recorded in head-fixed larvae appeared to form a continuum rather than a series of discrete group of movement types. Although previous studies have described discrete categories of movements,^4^ there are two reasons why some continuity is expected. The firing rate of reticulospinal neurons gradually influences the kinematic parameters of tail movements, such as the amount of turn for MiV1 neurons^5^ or the tail-beat frequency for nMLF.^10^ Such populations of neurons are thought to modulate continuously the trajectories.^11^

In order to obtain a parsimonious representation of this continuum of locomotor actions, I trained a supervised classifier to identify 5 different categories of movements: scoot, asymmetrical scoot, routine turn, C bend and burst. Tail bouts were correctly classified with 82% chance while errors in the classification occurred mostly between similar categories. Although previous studies have performed categorization of behavior in free-swimming conditions, our method does not rely on the analysis of the generated trajectories and enabled me to analyze the behavior during functional imaging and to produce a fine description of the relation between neuronal activity and locomotion.

## Supplementary Methods

### Feature Extraction

To compute the feature of the curvature matrix of a tail bout, I concatenated features from the time series of the curvature along the tail’s length. Each time series represented the curvature at a specific location of the tail, I computed statistical moments up to the 4th order, maximum, minimum, number of peaks, number of zero crossing and Fourier coefficients. After this, a PCA was applied and only the subspace corresponding to the 15 largest components was kept.

### Dynamic Time Warping

The DTW algorithm has earned its popularity by being an efficient similarity measurement for time series in areas such as data mining,^12^ gesture recognition^13^ or speech processing.^14^ DTW reduces the distance between the time series by warping the time axis. Given two tail bouts for which the tail deflection index were *X* = (*x*_1_,*x*_2_,…,*x*_*N*_) and *Y* = (*y*_1_,*y*_2_,…,*y*_*N*_) with *N* = 30 corresponding to a bout duration T=150ms and a sampling frequency Fs=200 Hz. DTW yield an optimal solution in *O*(*N*^2^) time. The algorithm starts by building the distance matrix *C* ∈ ℝ^*N*\**N*^ representing all pairwise distances between X and Y (Fig. 6.A). Once the cost matrix is built, the algorithms aims at finding the lowest energy path in the matrix (Fig. 6.A). The alignment path built by DTW was a sequence of points *p* = (*p*_1_,*p*_2_,…,*p*_K_) with *p*_l_ = (*n*_i_,*m*_j_) ∈ [1: *N*]^2^ for l ∈ [1: *K*] which satisfied the following criteria:

- Boundary condition: *p*_1_ = (1,1) and *p*_*K*_ = (*N,N*). The starting and ending points of the path must be the first and last points of the sequences.
- Monotonicity condition: *n*_1_ < *n*_2_ < … < *n*_*K*_ and *m*_1_ < *m*_2_ < … < *m*_*K*_. This condition preserves the time-ordering of the points.
- Step-size condition: *p*_*l*+1_ – *p*_l_ ∈ {(1,1), (1,0), (0,1)}. This criteria limits the shifts in time from large jumps.

**Figure 6.**
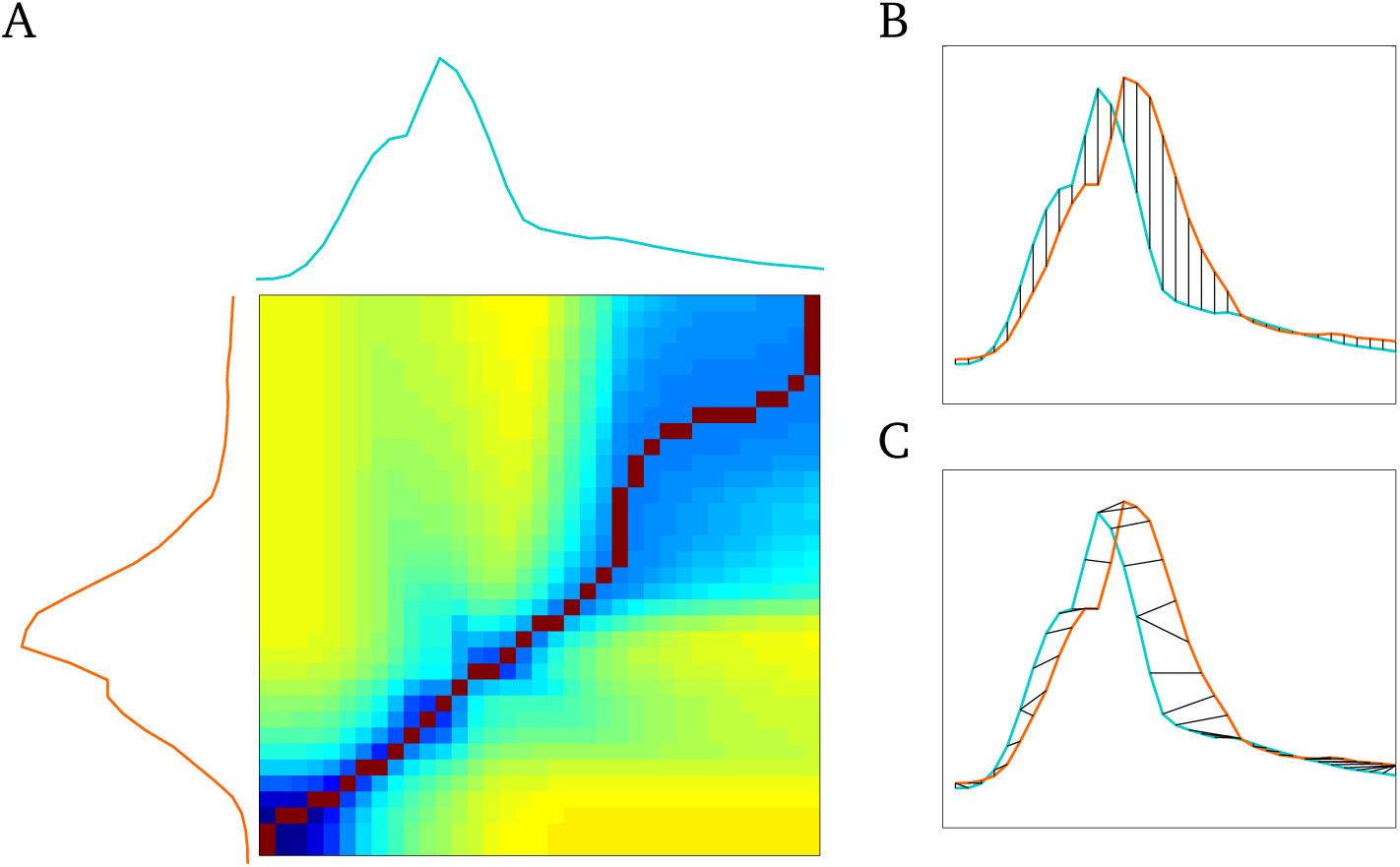
Alignment of tail deflection using DTW. (**A**) Cost matrix for the time series of two tail deflection indexes in cyan and orange. The optimal alignment path is shown in red. (**B**) An example of measurement of the euclidean distance for the two time series of tail deflection in (A). The comparison point by point is unable to recover the similarity between the two time series because of the small temporal offset. (**C**) Optimal time alignment of the two sequences is shown in black.

DTW uses dynamic programming to find the path of minimal cost *p*^*^:

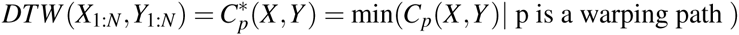

The optimal path *p*^*^ is computed in reverse order of the index, starting with *p*_*L*_ = (*N,N*). Supposing that *p*_l_ = (*n, m*) has been computed. In the case where (*n,m*) = (1,1), one must have l = 1 and the algorithm is complete. Otherwise,

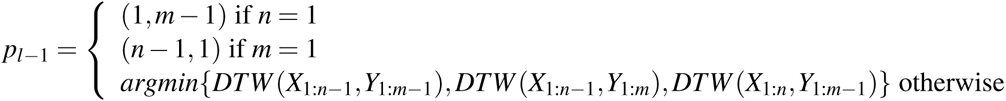

It should be noted that DTW is a similarity measure or a pseudo-metric. Unlike the euclidean distance, it does not verify triangular inequality required for metric.

### Fuzzy k-nearest neighbor classification (FKNN)

The k-nearest neighbor is the simplest non-parametric classification method.^15^ A class is assigned according to the most common class among its k-nearest neighbors. In FKNN, a membership level u_i_ is assigned to a tail movement x using the following formulation:

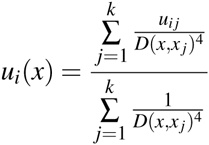

where *i* = 1,2, …,5 corresponds respectively to the category: Scoot, Asymmetric Scoot, Routine Turn, C Bend, Burst. *x*_*j*_ are the tail bouts in the library of user-defined labeled tail bouts (containing ~ 800 tail bouts). The distance D is the DTW distance. The factor 4 is used to weight the similarities and was found to minimize the error in classification. We only consider the *K* = 10 first neighbors. The *u*_*ij*_ terms represent the membership degree of the movement *x*_*j*_ from the training set to the class *i*. They are defined as:

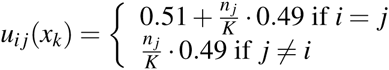

The value *n*_*j*_ is the number of neighbors which belong to the *j*-th class. Thus even tail movements that where manually labeled have a fuzzy membership to account for error during the manual labeling. Note that the sum of memberships *u*_*i*_ is equal to 1 making the interpretation as a probability straightforward. The direction (or bias) of movements belonging to asymmetric categories: Asymmetric Scoot, Routine Turn and C Bend was defined by the sign of 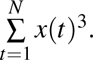.

### t-Distributed Stochastic Neighbor Embedding

Dimensionality reduction techniques such as PCA, multi-dimensional scaling or Isomap minimize the deformation of large distances.^16,17^ Similarity measurements such as DTW are more relevant at the level of the local neighborhood. I chose t-SNE for visualization because it aims at minimizing local distortions. For t-SNE, the conserved invariants are related to the Markov transition probabilities of a random walk performed on the dataset. Specifically, it defines the probability of transition from a tail movement xi to another *x*_*j*_, *p*_*j* *i*_, to be proportional to a Gaussian kernel of the distance between them:

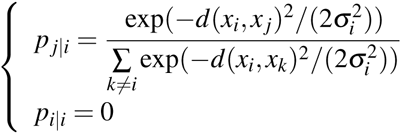

Each σ_*i*_ is set such that all points have the same transition entropy: 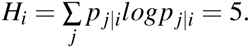 This parameter can be interpreted as controlling the number of neighbors considered, similar to *K* in the FKNN. Then, t-SNE places the points in the 2D euclidean space where the transition probabilities *q*_*j|i*_ are as similar to *p*_*j|i*_ as possible. *q*_*j|i*_ is chosen, for technical reason, to be proportional to a Cauchy (or Student-t) kernel of the distance between points in the 2D space. This algorithm results in an embedding that minimizes local distortions. *p*_*j|i*_ with small values corresponding to dissimilar tail movements will impose little constraints on the embedding. The complexity of the method in *O*(*N*^2^) restricts its applications to datasets that contain no more 10 000 points. I used the Matlab implementation provided by the authors of the algorithm.

## Acknowledgements

This work was supported by EraSysBio+ Zebrain, ERC stg 243106, ANR-10-LABX-54 MEMO LIFE, ANR-11-IDEX-0001-02 and the PSL Research University. A.J acknowledges support from the *Fondation pour la Recherche Médicale and the ENS Cachan*.

## Author contributions statement

A.J conceived the experiments, performed behavioral assay, analyzed the data, and built the setup and software. G.S designed the project and contributed to manuscript writing.

## Additional information

The authors declare no competing financial interests.

